# Generative lesion pattern decomposition of cognitive impairment after stroke

**DOI:** 10.1101/2020.08.22.262873

**Authors:** Anna K. Bonkhoff, Jae-Sung Lim, Hee-Joon Bae, Nick A. Weaver, Hugo J. Kuijf, J. Matthijs Biesbroek, Natalia S. Rost, Danilo Bzdok

## Abstract

Cognitive impairment is a frequent and disabling sequela of stroke. There is however incomplete understanding of how lesion topographies in the left and right cerebral hemisphere brain interact to cause distinct cognitive deficits. We integrated machine learning and Bayesian hierarchical modeling to enable hemisphere-aware analysis of 1080 subacute ischemic stroke patients with deep profiling ∼3 months after stroke. We show relevance of the left hemisphere in the prediction of language and memory assessments, while global cognitive impairments were equally well predicted by lesion topographies from both sides. Damage to the hippocampal and occipital regions on the left were particularly informative about lost naming and memory function. Global cognitive impairment was predominantly linked to lesioned tissue in supramarginal and angular gyrus, the postcentral gyrus as well as the lateral occipital and opercular cortices of the left hemisphere. Hence, our analysis strategy uncovered that lesion patterns with unique hemispheric distributions are characteristic of how cognitive capacity is lost due to ischemic brain tissue damage.

## Introduction

In the US, somebody experiences a stroke every 40 seconds (Benjamin *et al*., 2017). Ensuing ischemic brain lesions do not only underly motor impairments, but also cause a continuum of cognitive impairments ranging from clinically silent to overtly disabling symptoms (Jokinen *et al*., 2015). Variability in cognitive consequences underscore the clinical importance of an early and accurate prediction of cognitive outcomes at an individual-patient level. Further, personalized prediction of long-term post-stroke cognitive recovery using stroke lesion topography may offer valuable cues for tailoring future therapies and rehabilitation strategies.

However, the precise structure-function correspondence of how stroke lesions are linked to distinct cognitive deficits will pave the way for more reliable predictions of clinical endpoints. For instance, thalamus and angular gyrus lesions relevant for cognitive performance are typically located in the language-dominant hemisphere. In lesion studies focused on picture naming, the tissue integrity of the left superior and anterior to posterior middle temporal gyrus as well as the left inferior parietal lobe was vital for sustained naming abilities (Damasio *et al*., 2004; Baldo *et al*., 2013). Further, limbic system regions, such as the hippocampus and amygdala, are well known to be implicated in memory functions in the intact human brain (Papez, 1937).

Many brain functions involve various lower and higher cognitive processes and are supported by regions spread across the brain’s grey matter. A key aspect of the spatial distribution of neurocognitive processes is the preferential lateralization to one hemisphere. Two of the potentially most consistent aspects of hemispheric functional lateralization, language (Broca, 1861; Wernicke, 1974) and attention (Corbetta and Shulman, 2002), may distinctly relate to the left versus right hemisphere. Indeed, language impairments occur more frequently and severely after left-and neglect after right-hemispheric strokes.

Previous stroke lesion studies have typically been conducted under some anatomical constraint to tame the amount of information in lesion volume and distribution (Vaidya *et al*., 2019). Hence, these language impairments and neglect are commonly studied specifically in the predominantly affected hemisphere (language impairment & left hemisphere: Bates *et al*., 2003; Baldo *et al*., 2013, neglect & right hemisphere: Vallar and Perani, 1986; Smith *et al*., 2013). For instance, one of the earliest studies on neglect examined overlay lesion maps of 110 patients with right-sided stroke (Vallar and Perani, 1986). Nonetheless, several studies suggest that acute neglect also affects more than 20% of left-hemispheric stroke patients (Ringman *et al*., 2004). In a similar vein, aphasia is mostly caused by lesions in the hemisphere contralateral to an individual’s dominant hand. However, there are cases of “crossed aphasia” (Coppens *et al*., 2002), implying language deficits due to lesions in the right hemisphere, *ipsilateral* to the dominant hand.

Therefore, in this proof-of-principle study, we sought to integrate machine learning and Bayesian hierarchical modeling to enable hemisphere-aware analysis of 1080 subacute ischemic stroke patients.

## Methods

### Participant recruitment

Using a prospective stroke registry database (Kim *et al*., 2015), we identified hospitalized stroke patients who were diagnosed with acute ischemic infarction based on diffusion-weighted magnetic resonance imaging (MRI) typically within 1 week after symptom onset between January 2007 and December 2018 in Hallym University Sacred Heart Hospital or Seoul National University Bundang Hospital, South Korea. Among them, a total of 1080 patients were selected based on the following criteria: (1) availability of brain magnetic resonance imaging (MRI) showing the acute symptomatic infarct(s) on diffusion-weighted imaging (DWI) and/or fluid-attenuated inversion recovery (FLAIR), (2) successful infarct segmentation and registration (see below), (3) no previous cortical infarcts, large subcortical infarcts (>15 mm) or hemorrhages (>10 mm) on MRI, and (4) availability of data on key demographics and neuropsychological assessment (the 60-min Korean-Vascular Cognitive Impairment Harmonization Standards-Neuropsychology Protocol, K-VCIHS-NP; Hachinski *et al*., 2006; Yu *et al*., 2013) within one year post-stroke. We excluded patients (1) whose MRI was inadequate for properly obtaining neuroimaging variables, (2) who had bilateral stroke, and (3) inability to undergo cognitive testing due to severe aphasia, as determined by the attending physician.

Stroke subjects provided informed written consent in accordance with the Declaration of Helsinki. The local institutional review boards approved the study protocol and gave waived consent requirements based on the retrospective nature of this study and minimal risk to participants.

### Characteristics of participant sample

Stroke patients underwent a comprehensive battery of neuropsychological tests ∼3 months after onset of acute stroke (median time post-stroke: 98 days; Yu *et al*., 2013). We here predominantly focused on three key assessments of post-stroke cognitive performance: Korean Mini-Mental State Examination (MMSE, total score; Folstein, Folstein and McHugh, 1975), language performance (total score, Korean short version of the Boston Naming Test (BN); Kaplan, Goodglass and Weintraub, 1983) and memory function (Immediate recall, Seoul Verbal Learning Test (SVL), the Korean equivalent to the California Verbal Learning Test; Yu *et al*., 2013). Performance in the MMSE reflects global cognition including the orientation to time and place, as well as calculation or language performance. The BN (Korean short version) is a clinical standardized test that measures the naming abilities of patients. The SVL, in turn, examined episodic memory performance and requires the auditory learning of a word-list and tests its memorization by an immediate recall task.

The cognitive performance prior to stroke onset was captured by the Informant Questionnaire on Cognitive Decline in the Elderly (IQCODE; Lee *et al*., 2005). This test provides a widely used and validated measurement instrument of cognitive decline in the ten years prior to stroke onset and relies on health-proxy reports. Additionally, available sociodemographic and clinical information included: age, sex, years of education and lesion volume (**Table 1)**. Continuous (non-binary) variables were put on a more comparable scale by normalization (MMSE, BN, SVL, lesion volume, age, education years).

**Table 1.** Patient characteristics.

|  | **Mean (95%-CI)** |
| --- | --- |
| **Age** | 67.4 ± 0.7 (range 31 – 94; median 69.0) |
| **Sex** | 57 % male, 43% female |
| **Clinical history of stroke** | 12.6 % |
| **Years of education** | 9.3 ± 0.3 (range 0 – 23; median 9) |
| **IQCODE** | 3.38 ± 0.04 (range 1.3 – 5.0; median 3.2) |
| **Mini-mental state examination (total score)** | 24.0 ± 0.4 (range 0 – 30; median 26) |
| **Boston naming test (total score)** | 10.1 ± 0.2 (range 0 – 15; median 11) |
| **Seoul verbal learning (immediate recall)** | 15.2 ± 0.4 (range 0 – 36; median 16) |

### Neuroimaging data preprocessing

Brain-imaging acquisition was typically performed within the first week after stroke. Brain scanning included structural axial T1, T2-weighted spin echo, FLAIR and DWI sequences (3.0T, Achieva scanner, Philips Healthcare, Netherlands, image dimensions: 182×218×182; c.f., supplementary materials for details). Stroke lesions were manually segmented on DWI or less frequently FLAIR images by experienced, trained investigators (A.K.K., G.A.) relying on in-house developed software based on MeVisLab (MeVis Medical Solutions AG, Bremen, Germany; Ritter *et al*., 2011). Lesion segmentations were subsequently checked and adapted by two experienced raters (N.A.W., J.M.B). Subsequently, images as well as corresponding lesion maps were linearly and non-linearly normalized to Montreal Neurological Institute (MNI-152) space employing the RegLSM image processing pipeline (public code: http://lsm.isi.uu.nl/; Weaver *et al*., 2019). Quality of normalization was rigorously controlled by an experienced rater (N.A.W.). If there were any visual differences between the original and registered lesion maps during quality control, normalized lesion maps were manually corrected.

To develop a fully probabilistic generative modeling framework specifically tailored to stroke populations, we combined: 1) unsupervised dimensionality reduction of lesion maps and 2) Bayesian hierarchical modeling to predict the outcome in cognitive performance tests. We here focused on quantitative analyses of the 3-months outcomes measured on the MMSE, Korean short version of the BN and the SVL.

### Data-driven exploration of anatomic lesion representations

Each lesion map to-be-analyzed featured 435,642 total voxels of 1 mm^3^ in grey matter. We sought to explore coherent topographical patterns that may be hidden in these high-dimensional brain scans and represent each patient’s lesion fingerprint in a simpler and more directly interpretable form. To this end, we first computed the lesion load within 54 parcels (108 for both hemispheres), composed of the Harvard-Oxford cortical atlas with 47 regions and subcortical atlas with 7 regions in each hemisphere. We first summarized the number of voxels affected per atlas-defined brain region. In doing so, we obtained 54 regional measures of lesion load per hemisphere in each participant. We log-transformed and concatenated the ensuing lesion load measures for the left and right hemisphere. We then applied non-negative matrix factorization (NMF; Lee and Seung, 1999) as a multivariate encoding strategy to identify unique combinations of spatially distributed region damage (c.f. supplementary material for further details and justifications). NMF achieved a low-rank approximation of the anatomical lesion data by partitioning the lesion maps into a matrix of latent factor representations **W**. These hidden topographical representations link the emerging spatial lesion pattern topography to each of the original anatomical brain regions. The matrix of latent factor loadings **H** indicates how relevant each emerging spatial lesion pattern is to describe a specific patient’s overall lesion distribution. The atomic representations hence decompose the actual lesion constellation in a given patient given by **V** = **WH**, with **V** being the local lesion summaries of p x n dimensions (p = number of brain regions and n = number of subjects). Accordingly, **W** and **H** have p x k and k x n dimensions, respectively, with k representing the number of latent spatial patterns.

NMF provided at least two key advantages: In contrast to classical lesion-symptom mapping(Bates *et al*., 2003), that considers the effect of one location at a time, each brain location could here belong to several latent lesion components of **W** to varying degrees. In this way, each location could contribute to the prediction of cognitive outcome through relative contributions of multiple components, each of which reflected extracted lesion archetypes distributed across the whole brain. In this way, it was more similar to multivariate-lesion symptom mapping approaches (e.g., Smith *et al*., 2013) and also alleviates the likelihood and amount of distorted functional localization due to functional dependence (Mah *et al*., 2014; Sperber, 2020). Additionally, the non-negativity of the segmented brain lesion information and the non-negativity constraint of the NMF model ensured the automated derivation of a parts-based representation. That is, each individual latent component **W**_**a**_ represented a unique and directly interpretable aspect of the overall topographical lesion pattern variation. The biologically more meaningful component representations issued by NMF are in stark contrast to latent representations learned by alternative matrix factorization algorithms. For instance, in principal component analysis (PCA), individual lesions would be recovered through convoluted additions and subtractions of several components, which is why the ensuing components would not have been as easily and intuitively interpretable.

### Predicting interindividual differences in cognitive outcomes

The NMF-derived expressions of topographical lesion atoms provided the neurobiological input into our Bayesian hierarchical model (Gelman and Hill, 2006) to explain interindividual differences in cognitive outcome scores. We opted for this generative, multilevel approach given our primary motivation to quantify the full probabilistic information in our model parameter estimates at several neurobiological organizational levels. In particular, the obtained uncertainty distributions therefore made explicit the confidence of the model for each hemisphere and each lesion atom in contributing to the successful prediction of a given cognitive outcome. We have designed a bespoke Bayesian hierarchical models dedicated for each cognitive outcome (i.e., MMSE, BN or SVL, **Table 1)**.

To directly examine possible differences in hemispheric predictive relevance in our data, the modeled generative process assumed a joint dispersion prior for all lesion atoms of each hemisphere. Therefore, the standard deviation priors for left and right hemisphere could capture the hemisphere-specific predictive contributes tiled across all candidate lesion atoms. Priors of left-and right-hemispheric standard deviations were additionally combined through a joint hyperprior to complement the hierarchical model structure. Furthermore, the model took into account several covariates, including age, age^2^, sex, age-sex interactions, education years, pre-morbid cognitive performance and total lesion volume. We could then inspect these covariates in conjunction with hemisphere or lesion atom contributions to concurrently capture the contribution of a lesion representation to the cognitive outcome while rendering explicit the influence of important demographic, sociodemographic and clinical factors (c.f. **Supplementary figure 1** for the full model specification).

Samples from the posterior distribution of the model parameters were drawn by the No U-Turn Sampler (NUTS), a kind of Monte Carlo Markov Chain algorithm (setting: draws=5000; Hoffman and Gelman, 2014). Posterior predictive checks were carried out after model estimation to evaluate the obtained predictive model. That is, we empirically assessed the simulated outcome predictions generated by our model solution to approximate external validation based on our patient sample.

Our Bayesian hierarchical approach facilitated the careful dissection of predictive relevance allocated to different levels of the model. For all outcome models, we first evaluated lateralization effects that were inferred from the left and right hemisphere posterior sigma distributions. In analogy to ANOVA applications, where the total variance of a specific outcome variable is partitioned into several sources of variance, we thus here focused on the proportion of the variance that was attributable to variation in all lesion atoms linked to one brain hemisphere.

In this way, we capitalized on the general hemispheric importance in predicting the outcome, yet de-emphasized directionality of these associations: “Did either one hemisphere, the left or right one, contribute more substantially to the accurate prediction of the outcome of interest?” Therefore, predictive relevance could originate from predictors of more (“more impaired”) or less deteriorated (“less affected”) cognitive impairment observed in our patient sample. Subsequently, we considered lateralization effects of specific lesion atoms and lastly reverted back the predictive relevance of lesion atoms (Bzdok *et al*., 2016) to the level of the anatomical brain regions for each of the cognitive outcomes.

### Code availability

Analyses were conducted in a Python 3.7 environment and predominantly relied on the packages nilearn and pymc3. Full code for reproducibility and reuse is available here: www.github.com/TO_BE_ADDED

## Results

### Characteristics of cognitive outcomes and patient sample

We relied on structural MRI data of 1080 ischemic stroke patients, typically obtained within one week after symptom onset, to predict cognitive outcomes ∼3 months post-stroke. Analyses were focused on the Korean Mini-Mental State Examination (MMSE, total score; Folstein, Folstein and McHugh, 1975), language performance (total score, Korean short version of the Boston Naming Test (BN); Kaplan, Goodglass and Weintraub, 1983) and memory function (Immediate recall, Seoul Verbal Learning Test (SVL), Yu *et al*., 2013). Patients scored 24 points on the MMSE on average. Out of 15 objects, patients correctly named on average 10 objects in the BN. In the SVL, patient could remember 16 out of 36 words on average. Moreover, patients had an average of 9.3±0.3 years of education and scored 3.38±0.04 points on the Informant Questionnaire on Cognitive Decline in the Elderly (IQCODE; Lee *et al*., 2005; **Table 1**).

### Anatomy of the extracted lesion atoms in stroke patient

Voxel-wise topographical overlap of the lesion distributions demonstrated an excellent whole-brain coverage across patients (**Figure 1**). In the majority of the patients, stroke affected either the left or right vascular territory of the middle cerebral artery (MCA). The maximum of lesioned tissue was localized in subcortical zones. Importantly, there was no difference in the number of lesioned voxels per patient between the left versus right hemisphere (two-sided t-test: *p*=0.81).

**Figure 1.**
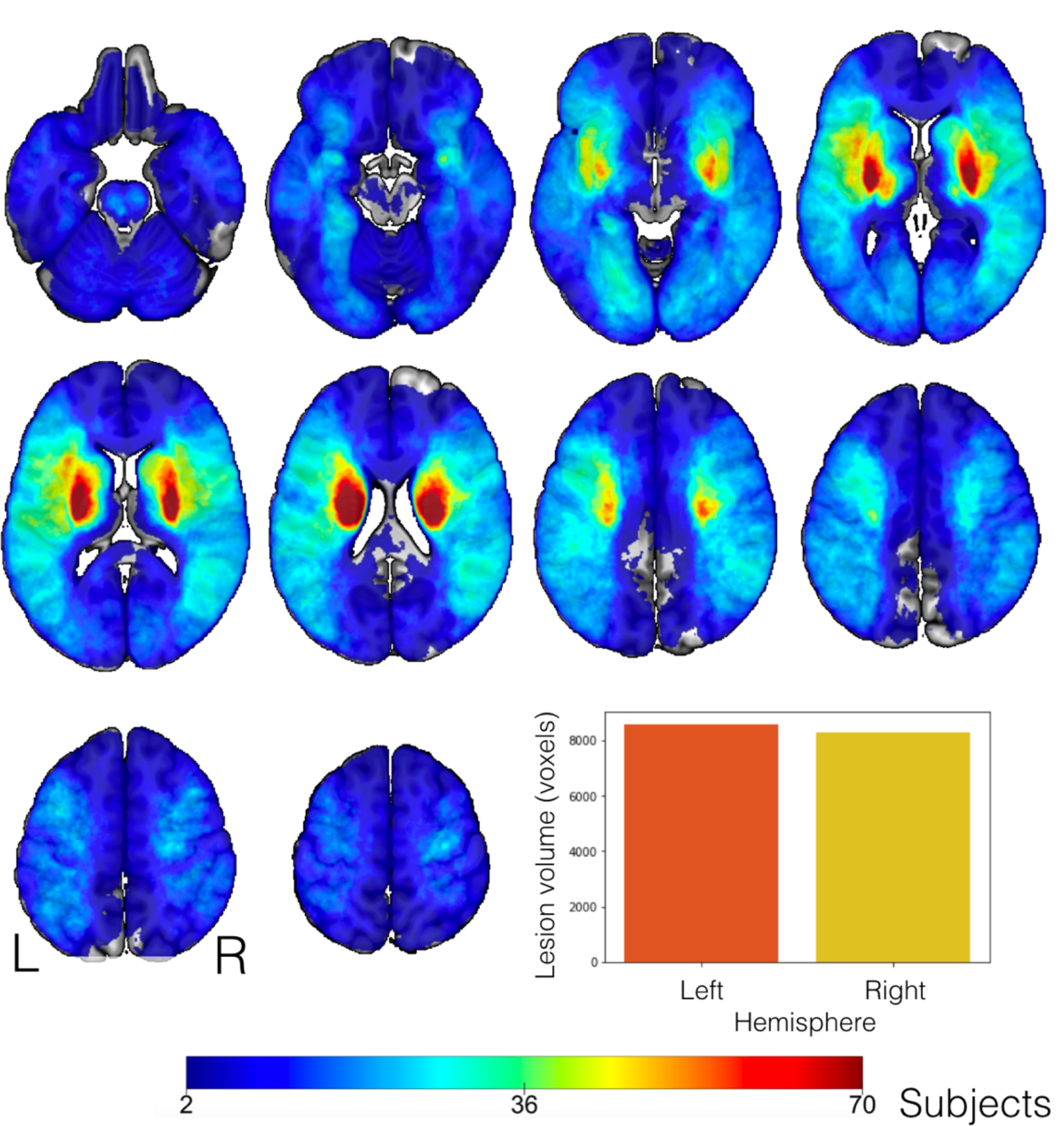
Topographical overlap of ischemic stroke lesions in 1401 patients in MNI reference space before the exclusion of patients with bilateral stroke. Tissue damage caused by neurovascular ischemic events affected predominantly subcortical regions, in line with previous reports (Corbetta *et al*., 2015). The strongest anatomical overlap in tissue lesions was found in the vascular territory supplied by the left and right middle cerebral arteria. We confirm excellent whole-brain coverage, with pronounced inter-individual heterogeneity in topographical distribution of tissue lesions. The overall lesion volume did not differ significantly between the left and right hemisphere in our patient cohort (t-test: *p*=0.81, bar plot lower right corner).

We summarized the initial high-dimensional lesion information at the voxel level in 108 cortical and subcortical brain regions for exploration of coherent hidden patterns of lesion topography. By these means, we uncovered a set of distinct topographical lesion configurations, *lesion atoms*, in a data-driven fashion based on non-negative matrix factorization (NMF; Lee and Seung, 1999). The spatial distribution of the lesion atoms corresponded to interpretable and biologically plausible components of stroke lesions and correlation patterns between lesions atoms were largely similar between the left and right hemisphere (**Figure 2**). Lesion atoms separately represented topographies in line with territories of arterial blood supply via the anterior (ACA, lesion atom 7), middle (MCA, cortical: 3, 5, 6, 10, subcortical: 1, 2, 8) and posterior (PCA, 9) cerebral artery territories.

**Figure 2.**
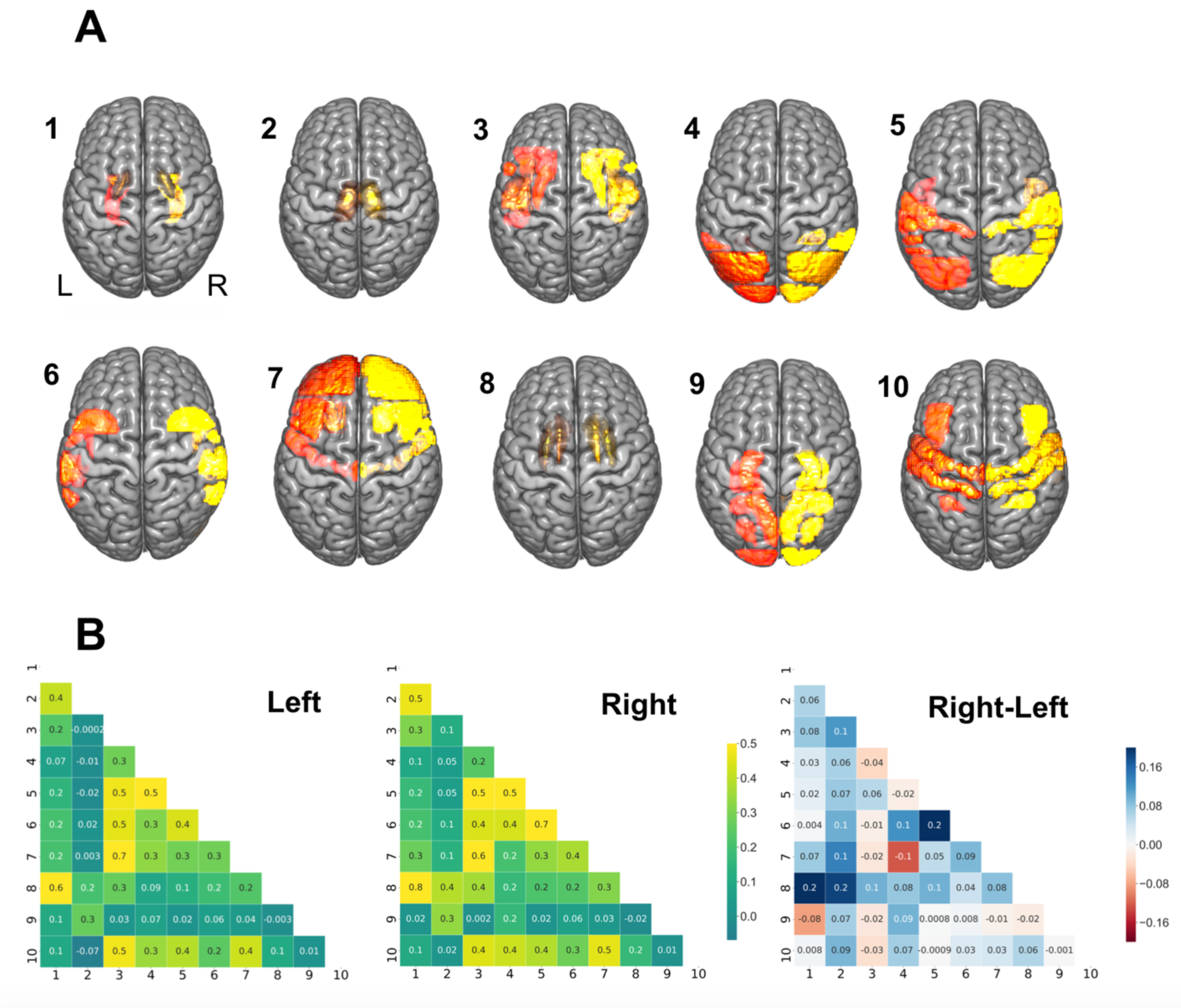
Lesion atoms of stroke patterns were derived by dimensionality-reducing pattern recognition. Voxel-wise lesions were summarized based on 108 brain region definitions (54 per hemisphere). The region-wise lesion measures were then further compressed into 10 essential lesion-pattern dimensions on each side by capitalizing on non-negative matrix factorization. **A**. The thus uncovered set of lesion atoms recapitulated several well-documented stroke lesion patterns and was further supported by biological plausibility of the lesion pattern anatomy. For instance, several lesion atoms delineated specific cortical and subcortical stroke patterns coherent with the anatomy of the middle cerebral artery (subcortical: lesion atom 1, 2, 8; cortical: 3, 5, 6, 10). Posterior cerebral artery strokes were appropriately captured by lesion atom 9. Lesion atom 7 depicted most topographies in line with strokes of the anterior as well as middle cerebral artery. In particular, lesion atom 7 (ACA & MCA) delineated the frontal pole, middle and inferior frontal gyrus, precentral gyrus, frontal orbital cortex, frontal opercular cortex. Lesion atom 9 (PCA) included regions of the inferior temporal gyrus, intracalcarine cortex, cingulate gyrus, precuneus and cuneal cortex, parahippocampal gyrus, lingual gyrus, temporal and occipital fusiform cortex, occipital pole, as well as hippocampus. The remaining lesion atoms (largely MCA) captured lesion variation in regions including the subcortical regions globus pallidus, putamen, caudate and thalamus, as well as insular and cortical MCA regions (middle & inferior frontal gyrus (pars opercularis), superior, middle & inferior temporal cortex, pre-and postcentral gyrus, opercular cortex (frontal, central, parietal), planum polare, Heschl’s gyrus, angular gyrus, supramarginal gyrus, superior parietal lobule, lateral occipital cortex and the occipital pole). Additionally, the topography of lesion atoms aligned with a certain vessel territory matched the frequency of its affection by ischemic stroke in clinical practice: we obtained more fine-grained subdivisions of stroke topographies due to middle cerebral artery strokes in view of the high frequency that this vascular supply territory was affected. *Left/right* hemispheric lesion atoms shown in orange/yellow. **B**. Lesion atoms exhibited only small hemispheric differences in pattern correlations. Lesion atoms in subcortical areas were particularly often simultaneously affected in patients and thus strongly mutually related in the left and right hemisphere (atoms 1, 2, 8). For lesions on the right side, putamen and caudate (atom 8) exhibited slightly more prominent correlations with pallidal (atom 1) and thalamic areas (*darker blue color*).

After extracting this set of coherent atomic patterns that together underlie stroke topographies, we investigated the differential effects of a) hemispheres, b) lesion atoms, and c) anatomical regions on the cognitive outcome after stroke.

### Hemisphere relevance for clinical outcome

We first examined possible hemisphere-specific effects on cognitive impairment in our stroke patients. We specifically investigated whether lateralized relevance varied across the three outcome scores. We relied on the posterior distribution of variance parameters at the hemisphere level that had inferred the magnitude of hemisphere-specific relevance for a given cognitive dimension. In case of MMSE, similarly distributed hemisphere posteriors suggested that the combination of all lesion atoms of one hemisphere were equally relevant for predicting global cognitive outcome (mean of the left hemisphere-level posterior distribution=0.0404, highest probability density interval of the left hemisphere-level posterior distribution covering 94%-certainty (HPDI)=0.0157-0.0668; right posterior mean=0.0402, HPDI=0.011-0.0713, **Figure 3, left plot**). While the left hemisphere was predictive of poorer MMSE performance, the right hemisphere was predictive of more preserved MMSE performance. In contrast, the left hemisphere showed a markedly higher predictive relevance for our analysis of BN and SVL (BN: left posterior mean=0.0575, HPDI=0.0125-0.111, right posterior mean=0.015, HPDI=0.0001-0.0361; SVL: left posterior mean=0.0852, HPDI=0.0331-0.152, right posterior mean=0.0272, HPDI=0.0001-0.0609, **Figure 3, middle and right plot**). That is, tissue lesions located in the left hemisphere contributed comparatively more to the predictability of the naming and verbal learning impairments. The left posterior distributions of the hemisphere variance parameter had a higher posterior mean as well as wider spread across all lesion atoms in case of BN and SVL. Hence, the hemisphere parameters in our multi-level Bayesian model were directly informative about the left-hemispheric dominance when explaining interindividual variation in cognitive performance profiles.

**Figure 3.**
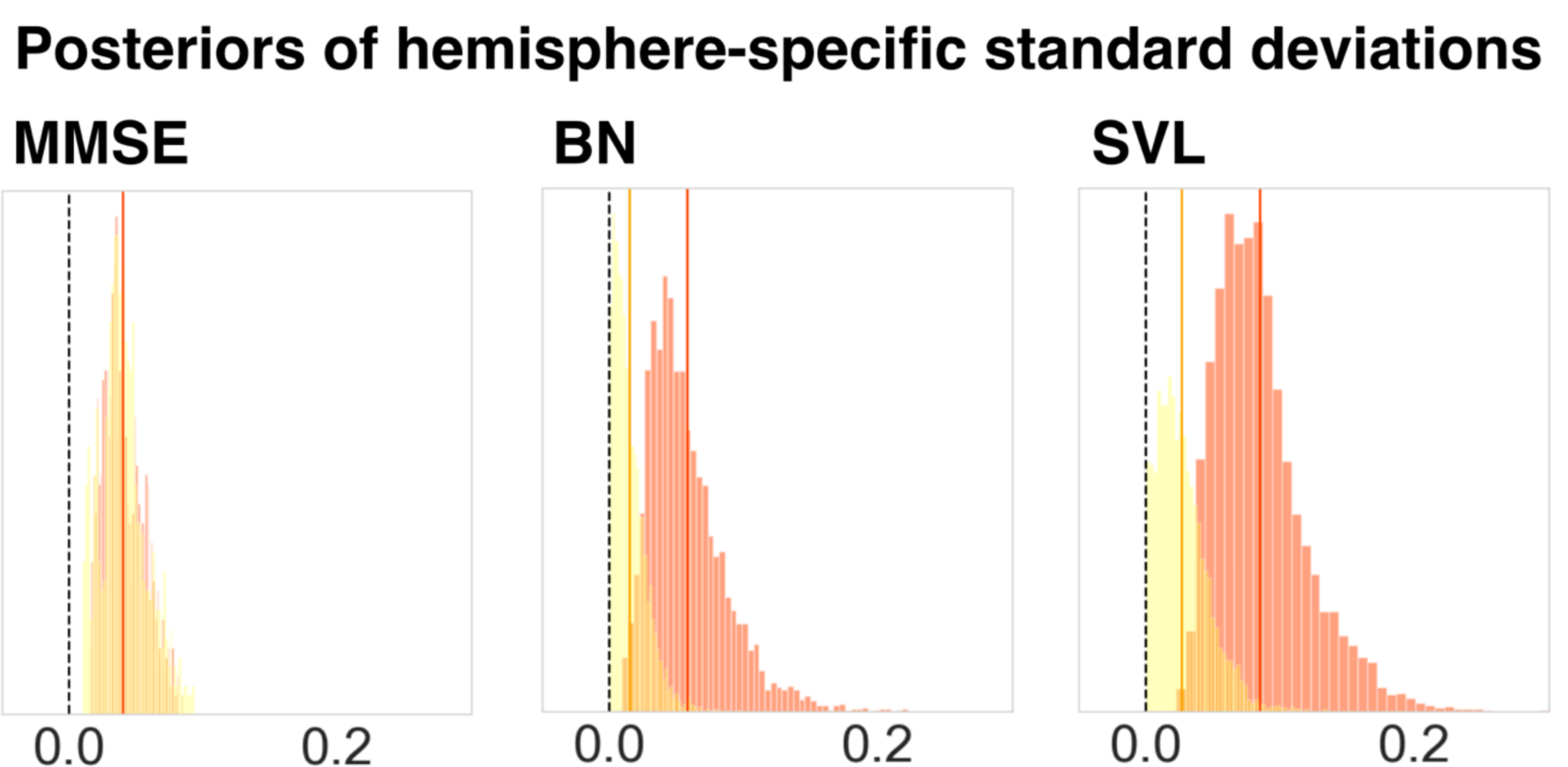
Hemispheric lateralization effects for predicting clinical outcomes in stroke patients. Displays the relevance of the posterior parameter distributions of the left and right brain. These were obtained through three Bayesian hierarchical models dedicated to predicting three cognitive outcomes. In case of the Mini-Mental State Examination (MMSE), often considered a measure of global performance tapping into various cognitive domains, lesions in the left and right brain contributed equally to prediction success. In contrast, lesions in the left brain were more relevant for single-patient predictions of cognitive outcomes in case of the Boston Naming (BN) and Seoul Verbal Learning Test (SVL). The shown posterior model parameters correspond to the upper hemisphere level of our Bayesian multi-level modeling strategy that uncovered the hemisphere-specific model certainty for each cognitive performance dimension. The revealed lateralization effects for BN and SVL highlight the left hemisphere (c.f., *vertical, dark orange lines*) that suggest a dominance of left-hemispheric lesion information in driving predictions for memory and verbal learning outcomes.

Furthermore, we observed characteristic inter-relations between these hemisphere-specific effects and important covariates, i.e., lesion volume, education attainment and pre-stroke cognitive performance, in the prediction of cognitive outcome (**Figure 4**).

**Figure 4.**
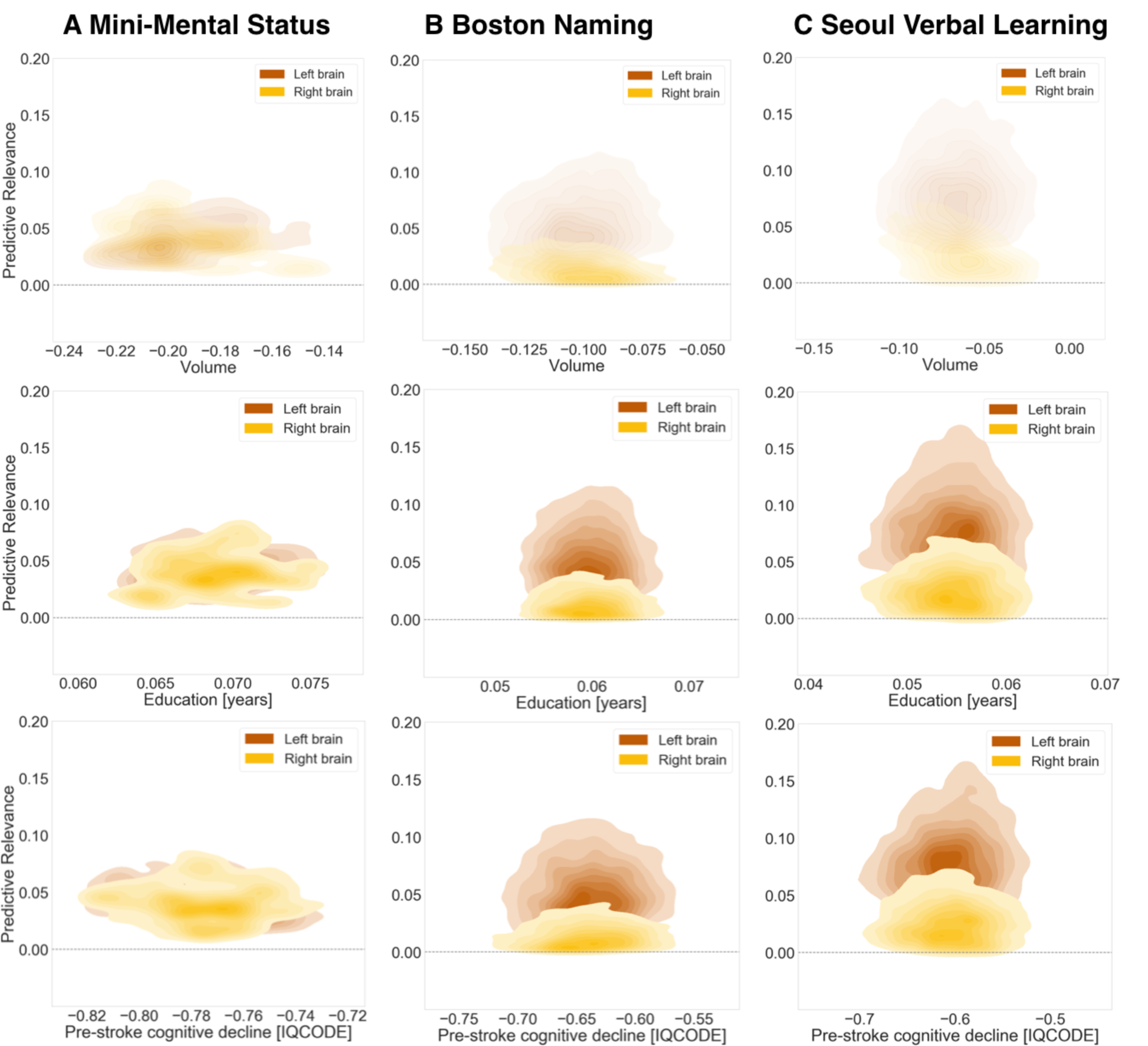
Hemispheric relevance and key covariates. Inter-relation between the relevance of lesion load in a given hemisphere (*y axes, left hemisphere: moccasin, right hemisphere: yellow*) and marginal posterior parameters of key covariates (*x axes)* in predicting cognitive performance. Lesion volume, as well as all three outcome variables were normalized to the same z score scale. Despite assumed important interactions (Adolphs, 2016), neither one of typically applied uni-or multivariate models represents an integrative approach to jointly study lesion location and further sociodemographic and clinical covariates. We here present such a joint analysis that is possible based on the combination of our large sample size and generative modelling framework. *Top row:* The influence of lesion volume on the cognitive outcome prediction varied across the three outcome scores: A one standard unit increase in lesion volume was sufficient to cause a 19% standard unit decrease in MMSE performance. Conversely, one standard unit increases in lesion volume led to only 7-10% standard unit decreases in BN and SVL performances. *Middle row:* Years of education had positive effects of comparable magnitude on all three outcome scores: An additional year of education predicted a 6-7% standard unit increase for MMSE, BN and SVL. *Bottom row:* One point more on the IQCODE scale, i.e. a higher pre-stroke cognitive decline, predicted a 60-64% drop in BN and SVL standard unit performance. In case of MMSE, on more IQCODE point resulted in an even higher decrease of 77% standard unit performance.

### Lesion atom relevance for clinical outcome

The level of lesion atoms in our hierarchical Bayesian model offered yet an additional opportunity to gain rich insights on lateralization effects linked to not only *entire* hemispheres, but specific lesion topographies *within* a hemisphere in the prediction of cognitive outcome. We thus direct our attention to the examination of individual lesion atoms that combined several brain regions. We inferred lateralization effects from non-overlapping posterior parameter distributions of left and right lesion atoms. Such non-overlapping distributions implied that a specific lesion atom in either the left or right hemisphere was reliably more predictive of a given cognitive impairment than the same lesion atom of the other hemisphere. Hemisphere-specific predictions showed relevant differences for a total of four out of ten lesion atoms considering all three target scores.

The first lesion atom was characterized by a pronounced left-lateralization of predictive relevance for BN and SVL (BN: hemispheric difference in the posterior mean=-0.0257, HPDI=-0.0487--0.00464; SVL: difference posterior mean=-0.0407, HPDI=-0.0708--0.00985). While the posterior distributions for the right-hemispheric first lesion atom did not meaningfully differ from zero, the left-hemispheric lesion atom distributions featured a pronounced deviation in the negative direction: Signaling that only lesions in the left lesion atom 1 regions were predictive of lost function in the BN and SVL (**Figure 6B&C, upper row**). The topography of the first lesion atom covered especially lesions of the globus pallidum and amygdala. Lesion atom 3 combined insular and opercular cortex, Heschl’s gyrus and the putamen and comprised lateralization effects for the MMSE, as well as SVL (MMSE: difference posterior mean=-0.0211, HPDI=-0.0524--0.00621; SVL: difference posterior mean=-0.134, HPDI=-0.227--0.0412). In case of SVL, the right-hemispheric posterior distribution once again did not diverge from zero and the left-hemispheric posterior distribution showed a marked shift into the negative direction. For the MMSE, the difference emerged due to lesions in the right hemisphere being slightly more predictive of preserved function and lesions in the left hemisphere being more predictive of lost function. For MMSE, we furthermore witnessed lateralization effects within lesion atom 4 that highlights lesions affecting supramarginal, angular and Heschl’s gyrus as well as lateral occipital cortex (difference posterior mean=-0.0651, HPDI=-0.124--0.00741). Lastly, BN and SVL led to marked left-lateralized predictive relevance in lesion atom 6 that comprised superior, middle and inferior temporal gyri, as well as lateral occipital cortex, planum temporale and amygdala (BN: difference posterior mean=-0.0746, HPDI=-0.147--0.00209; SVL: difference posterior mean=-0.0941, HPDI=-0.182--0.0115).

### Region relevance for clinical outcome

Coherent with our hemisphere-specific findings, we uncovered comparable predictive relevance at the brain-region level of the same estimated Bayesian model. Regarding MMSE, lesioned regions on the left were generally more predictive of lost function (negative parameter weights), while lesioned regions on the right were more predictive of preserved global cognitive functions (positive parameter weights). Explained variance, as determined via predictive posterior checks, for the MMSE model was R^2^=54.4% (**Figure 5B, left panel**).

**Figure 5.**
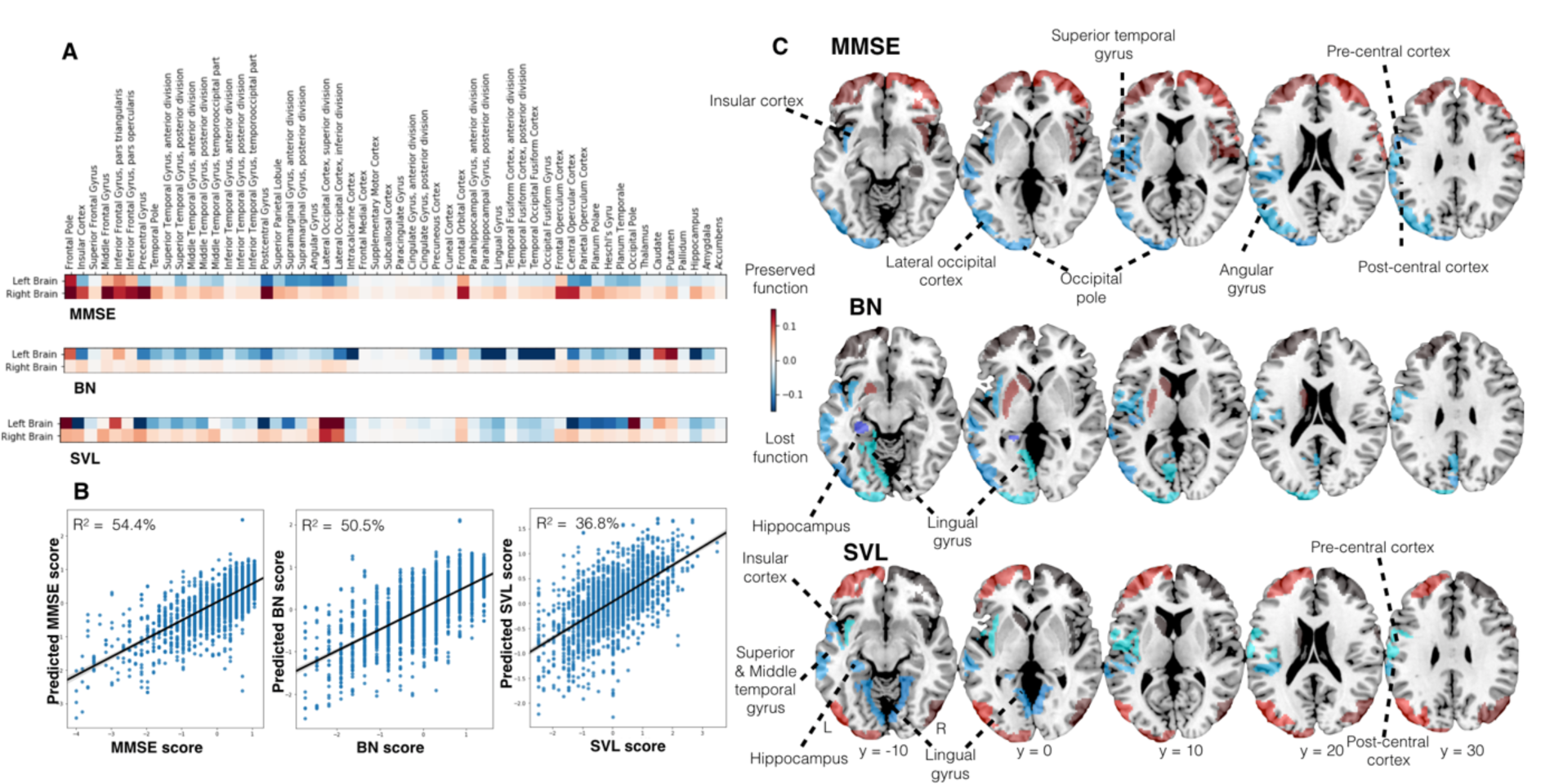
Cognitive impairments caused by stroke were predicted by unique lesion patterns. **A**. Heatmaps indicate the associations of each anatomical region (Harvard-Oxford atlas) with lost (*blue color*) or preserved (*red color*) clinical outcomes in patients. The Mini-Mental State Examination (MMSE) stuck out amongst all three scores as it featured predictive relevances that were equally prominent in the left and right hemisphere. However, these relevances were mainly indicative of preserved function (*red*) in the right hemisphere and of lost function on the left hemisphere (*blue*). The Boston Naming (BN) and Seoul Verbal Learning Test (SVL) were both characterized by pronounced left-hemispheric predictive relevances. These two tests differed in the exact region-wise distribution of the predictive relevances: Lesions predictive of poorer naming function primarily extended from the mediotemporal hippocampal to further occipital areas, while verbal learning impairments were better predicted by lesions affecting opercular and insular cortices, as well as planum polare and temporale. **B**. Prediction accuracy (R^2^ estimated via posterior predictive checks) for the three cognitive scores along with scatterplots of actual (*x axis*) and predicted (*y axis*) cognitive performances. The MMSE model achieved the highest R^2^ prediction performance (54.4%). Given its 50.5% explained variance, the BN model reached a slightly lower score. The explained variance of the SVL model totaled 36.8%. **C**. Brain renderings of the brain region-wise predictive relevance, described more in detail under **A**.

**Figure 6.**
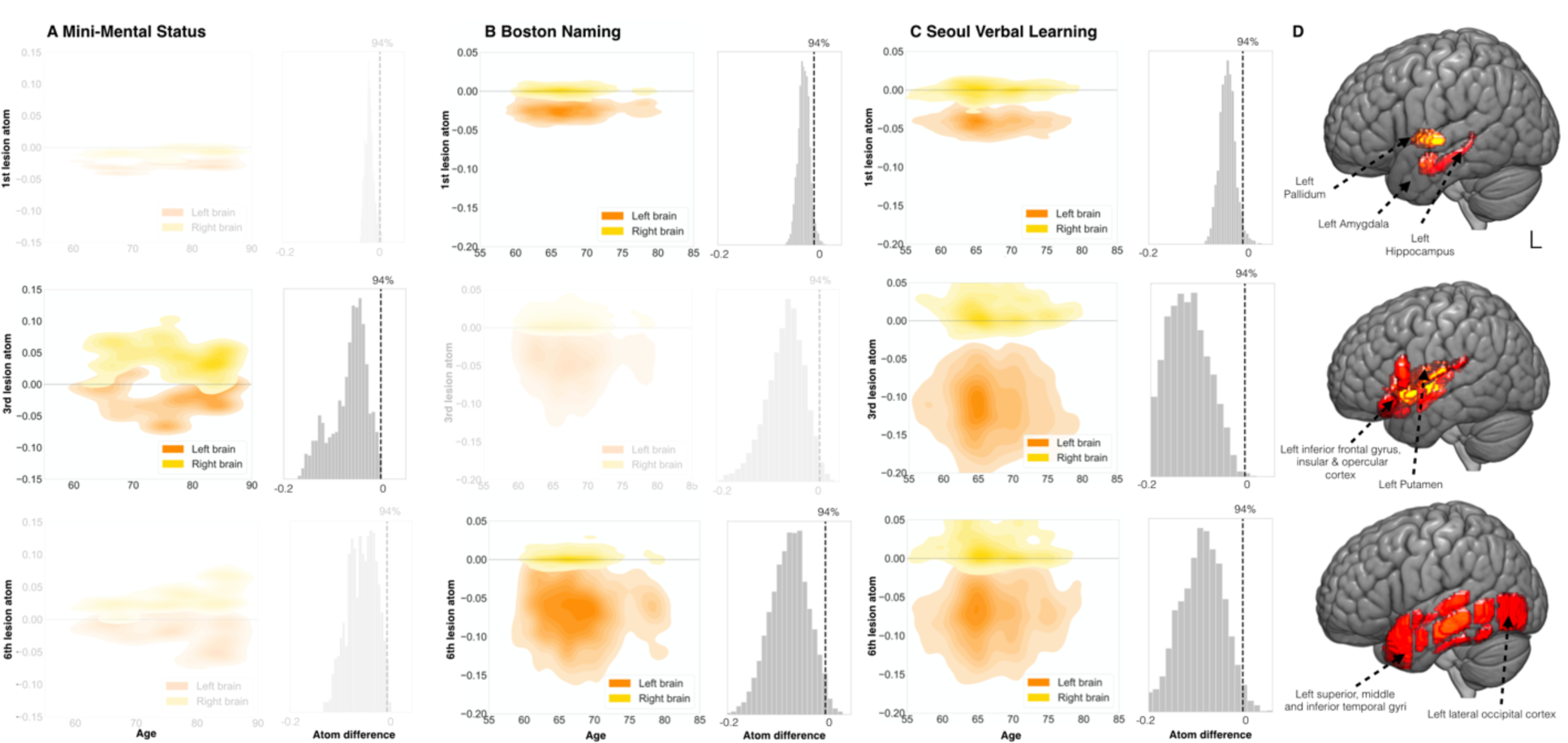
Specific lesion atoms show strong lateralization. **A-C**. Among all candidate lesion atoms, three distributed lesion patterns exerted pronounced hemispheric differences in predictive relevance to forecast future cognitive performance in at least two of the clinical outcomes. The Seoul Verbal Learning Test (SVL) was characterized by most lateralization effects of three lesion atoms. In contrast, lateralization effects were only observed for two lesion atoms for the Mini-Mental State Examination (MMSE) and Boston Naming Test (BN). Generally, we derived lateralization effects for lesion atoms by subtracting the marginal posterior distribution of one hemisphere from the marginal posterior distribution of the other hemisphere. We assumed a substantial lateralization when the distribution of the difference did not overlap with zero. In MMSE, hemispheric differences arose from right-hemispheric positive and left-hemispheric negative predictive relevances (positive: predictive of preserved function, negative: predictive of lost function). Left lateralization for BN and SVL originated from the difference between negative left-hemispheric and neutral right-hemispheric predictive relevances. Right columns: Differences between left-and right-hemispheric lesion atom-wise predictive relevances. Figures are shown slightly transparent, in case of a non-defensible difference and thus absent lateralization (0 inside of 94% highest-posterior density interval). Left columns: Plots visualize the joint density for combinations of parameter weights for age and each of the three lesion atoms. Age is plotted on the x axis, weights of lesion atoms on the y axis. Left hemispheric lesion atom weights are shown in orange, right hemispheric ones are shown in yellow. **D**. The four lateralized lesion atoms predominantly represented subcortical structures (putamen, globus pallidus, amygdala, hippocampus) as well as insular and temporal lobe areas.

Among the 108 total regions for both hemispheres, top regions informative for the prediction of lost MMSE function were the supramarginal and angular gyrus, the postcentral gyrus as well as the lateral occipital and opercular cortices. To a lesser degree, the insular cortex, precentral cortex and superior and middle temporal gyri contributed to the prediction of lost function as well (**Figure 5A&C, upper rows**). Crucially, all of these areas were located in the left hemisphere. Lesions located in frontal areas of both hemispheres as well as right-hemispheric insular and pre-and postcentral lesions were predictive of comparably more preserved function.

Regarding BN, predictive lesion patterns were predominantly isolated in the left hemisphere. The full model explained R^2^=50.5% of the variance in the BN outcome (**Figure 5B, middle panel**). The extent of naming impairment was particularly predicted by lesions extending from the mediotemporal hippocampal to further occipital areas. Moreover, extended parts of the temporal gyrus, especially superior and middle temporal gyri, demonstrated predictive relevance of lost function (**Figure 5A&C, middle rows**).

Regarding SVL, predictive lesion patterns resembled those of the BN to a great extent, however with a slightly varied weighting: left-hemispheric lesions had a more pronounced magnitude of predictive relevance. Posterior predictive tests indicated an explained variance of R^2^=36.8% (**Figure 5B, right panel**). Left-hemispheric opercular and insular cortices, as well as planum polare and temporale predicted deteriorated SVL function. In contrast to the BN test, hippocampal as well as occipital lesions of *both* hemispheres were predictive of lost function in case of SVL (**Figure 5A&C, lower rows**).

## Discussion

The majority of unique constellations of brain tissue damage show some extent of specific cognitive impairments (Jokinen *et al*., 2015). We here demonstrate the value of Bayesian hierarchical frameworks to explicitly model several biologically meaningful levels of stroke pathology to explain interindividual differences in cognitive outcomes. We automatically distilled plausible cortical/subcortical lesion patterns from a large multisite sample of 1080 stroke patients. These lesion configurations, naturally recurring motifs across patients, faithfully revealed coherent lesion atoms for cortical and subcortical MCA infarction, as well as lesion patterns recapitulating the supply territories of the ACA and PCA.

Our generative modeling tactic enabled the conjoint appreciation of predictive lesion features at the level of brain hemispheres, spatially distributed lesion atoms and individual brain regions. At the hemisphere level of our hierarchical model, we confirmed substantial lateralization effects in the left brain for naming and verbal learning impairments after stroke. For more global measures of cognitive performance, quantified by the MMSE, both hemispheres turned out to be equally informative about the patients’ clinical profiles. Importantly, the left hemisphere was predictive of comparatively more deteriorated performance, while the right hemisphere predicted less deterioration. At the level of lesion patterns distributed across regions, we found lateralization effects on the left for patterns capturing damage in the inferior frontal gyrus, insular and opercular cortex, the temporal lobe as well as the basal ganglia. At the level of single brain regions, memory impairment predictions were especially supported by left-hemispheric PCA-supplied brain regions, which involved the damage to the left hippocampus.

Our approach departs from classical lesion-symptom mapping endeavors in several ways. Many previous stroke imaging research pursued approaches that explore the effect of one brain location at a time (Bates *et al*., 2003). The common unit of observation in stroke brain-imaging research was thus the single brain voxel. Analyzing brain locations one after another however renders this classical approach less biologically meaningful by omitting lesion interactions between brain locations, which can be spatially distant. Our proof-of-principle approach simultaneously appreciates effects in (and interrelations between) local regions, their spatially distributed lesion constellations and hemispheric idiosyncrasies in a probabilistic multilevel framework for clinical outcome prediction.

Our main focus was on lateralization effects linked to cognitive performances. From a broader perspective, lateralization is a particularly well-established clinical feature for two frequent cognitive impairments after stroke. Each of these conditions is seen in at least 40% of common acute stroke patients: spatial neglect (Ringman *et al*., 2004) and aphasia (Flowers *et al*., 2016). In numerous cases, previous stroke studies cannot truly evaluate such effects due to a priori selection of anatomical regions of interest in either the left or right hemisphere (Vaidya *et al*., 2019). Furthermore, many lesion studies threshold their segmented lesion maps to focus on brain regions that are affected in more than five to ten percent of patients at hand (Smith *et al*., 2013; Zhang *et al*., 2014). This common practice also often amounts to confining analysis and interpretation to regions from one half of the brain.

To complement these previous research approaches to stroke, we aimed to avoid restricting analyses to one hemisphere or measuring voxel-wise lesion load. Instead, we established a principled analysis framework to capture hemispheric predictive relevance as a function of the cognitive outcome at hand in a dedicated probabilistic parameter estimate. In view of this single estimate, this quantification of hemispheric relevance enabled us to put lateralization-focus interpretations on a firmer basis. In so doing, we witnessed strong left-lateralization for BN and SVL. Both of these tests target the evaluation of memory functions, yet also heavily rely on language functions (Kaplan, Goodglass and Weintraub, 1983; Yu *et al*., 2013). Thus, the observed left-lateralization may well reflect these language functions’ main location in our subjects’ left hemispheres. Conversely, we did not find any evidence of hemispheric predominance, yet rather equally distributed predictive relevance across both hemispheres for the MMSE. This balanced contribution of both hemispheres in MMSE resulted from left-hemispheric lesions being predictive of more deteriorated test performance, while right-hemispheric lesions were predictive of more preserved function. Therefore, the relation between the left hemisphere and test performance resembled the one extracted for BN and SVL. Given that we focused on predictability and not causality, our finding does not necessarily imply an improved test performance in case of right hemispheric lesions, yet merely a consistently less deteriorated performance compared to patients with left hemispheric lesions. Also, the prediction of more preserved MMSE performance for lesions in the right hemisphere might be erroneously caused by the MMSE’s insensitivity to cognitive deficits after right-hemispheric lesions, as frequently reported in the literature (Grace *et al*., 1995; Dick *et al*., 1984; Kupke, Revis and Gantner, 1993).

While we found left-lateralization for cortical regions classically known to subserve lateralized language function, we also discovered indicators of left hemisphere-specific effects in subcortical regions. More precisely, these effects of lateralization highlighted especially the pallidum and putamen.

Aphasia is traditionally viewed as a deficit of the cortical mantle. Nonetheless, speculations about an involvement of the basal ganglia is going back to the late 19^th^ century (Wernicke, 1974; Broadbent, 1872; Marie, 1906). The exact role of non-thalamic subcortical areas in language functions remains incompletely understood, despite various lesion-symptom and functional neuroimaging studies in small cohorts. Characterizations of subcortical aphasias are highly heterogeneous and structure-function correspondences were repeatedly found to be weak (Naeser *et al*., 1982; Nadeau and Crosson, 1997; Radanovic, 2017). Our lesion findings are coherent with the interpretation of an implication of left putamen and globus pallus in healthy language function. Furthermore, selective pallidal lesions have been well characterized in neurosurgical case studies of Parkinson patients. Several investigators have described declines in verbal learning and memory functions specifically after left-sided pallidotomy. For instance, Lang and colleagues (1997) noted word finding impairments in 69% of patients after left-sided pallidal lesions, yet only in 25% of patients with such lesions on the right (Lang *et al*., 1997). Similarly, further studies suggested a higher verbal memory impairment after left-than after right-sided pallidotomy (Riordan, Flashman and Roberts, 1997; Crowe *et al*., 1998). However, the collection of these studies relies on less than 70 total patients. Moreover, analyses were previously restricted to the evaluation of anatomically very circumscribed lesions. In our approach, we extracted lesion atom-wise probabilistic hemispheric effects in 1080 patients while accounting for all other extracted lesion atoms covering regions across the entire brain. By these means, we can add convincing evidence to previously small and locally restricted studies.

Leveraging our novel, hemisphere-aware modeling approach, we could also reliably confirm and substantially expand previous reports on certain individual brain regions: Across all examined scores, we found predictive contributions to function loss for the superior, middle and inferior temporal gyrus, as well as insular and angular regions on the left side of the brain. These regions were also outlined in previous lesion-symptom studies of global cognitive and naming functions (Damasio *et al*., 2004; Baldo *et al*., 2013; Munsch *et al*., 2016; Zhao *et al*., 2018).

Most previous work presuming lateralization has focused on language or attentional impairments related to neglect. Due to their anatomical localization, these impairments closely relate to MCA strokes. That is, strokes in the most commonly affected vascular territory (Bogousslavsky, Van Melle and Regli, 1988). Adding to these previous efforts, we here also identified predictive contributions of left-hemispheric lesions in the less commonly affected vascular supply territory of the PCA. The here uncovered brain regions exerted negative effects on naming and verbal memory functions and were collectively represented by one of our automatically extracted lesion atoms. The left hippocampus emerged among the implicated regions in this specific lesion atom, which matched its frequent blood supply by the posterior cerebral artery (Erdem, Yaşargil and Roth, 1993).

A few earlier observations in stroke patients exist that have hinted at pronounced cognitive impairments after left-sided stroke lesions due to PCA occlusions (Benson, Marsden and Meadows, 1974; Renzi, Zambolin and Crisi, 1987; Szabo *et al*., 2009). More specifically, in neuropsychological testing, patients with PCA stroke performed particularly poorly in naming tasks and verbal memory tests (Benson, Marsden and Meadows, 1974; Renzi, Zambolin and Crisi, 1987; Szabo *et al*., 2009). A study that contrasted 20 left-and right-hemispheric PCA-stroke patients reported no differences in MMSE performance. However, the investigators noted significantly lower performances of patients with left-versus right-hemispheric stroke in two verbal learning tests (Szabo *et al*., 2009). Importantly, all patients considered in this study had lesions exclusively in the PCA-territory and all involved hippocampal regions.

Our study strengthens this scarce evidence (based on <50 total patients) by carefully quantifying the extent of cognitive impairments as a general consequence of brain infarcts extending to the PCA territory. Since we did not detect any comparable PCA-related lesion patterns predictive of MMSE performance, our results are coherent with the conclusion that lesions in left PCA-supplied regions have a particularly deleterious effect on naming and verbal learning functions. In fact, these impairments might primarily relate to hippocampal lesions, as captured in our lesion atom with PCA coverage.

Only recently, it was asserted that interindividual differences in several cognitive domains, such as verbal learning and global cognition, could arise from natural variability in vascular supply anatomy of the hippocampus (Perosa *et al*., 2020). These authors argued that a mixed supply, combining vessels from anterior as well as posterior cerebral artery, was more protective of cognitive impairment than a solely PCA-dependent supply. This study relied on data from healthy older adults as well as patients with cerebral small vessel disease, which promises future translation to stroke patients and their cognitive recovery.

## Conclusion

In conclusion, we have introduced an analytical framework that pools information across the entire brain and can therefore simultaneously integrate lesion facets on varying types and forms. The resulting multi-level estimates of predictive relevance may eventually finesse the probabilistic prediction of cognitive impairment at the single patient level.

## Acknowledgements

We gratefully acknowledge Angelina K. Kancheva and Gözdem Arikan for their help with performing the manual infarct segmentations.

## Author contributions

D.B., A.K.B.: Conception and design of study; D.B., A.K.B., J.S.L., H.J.B., N.A.W., H.J.K., J.M.B: Acquisition and analysis of data; D.B., A.K.B., N.S.R.: Drafting of significant portions of the manuscripts and figures; all authors: editing and approving the text.

## Supplementary material

### Neuroimaging data

**Seoul National University Bundang Hospital**

|  |  |
| --- | --- |
| **FLAIR** | repetition time: 11,000 ms; echo time: 125 ms; inversion time: 2,800 ms; slice thickness_ 5 mm; intersection gap: 1 mm; matrix: 512 x 512; flip angle 90 degree |
| **DWI** | EPI-spin echo sequence; repetition time: 5,000 ms; echo time: 50 ms; diffusion b-value: 1,000; slice thickness: 5 mm; intersection gap: 1 mm; matrix: 256 x 256; flip angle 90 degree |

**Hallym University Sacred Heart Hospital**

|  |  |
| --- | --- |
| **FLAIR** | repetition time: 11,000 ms; echo time: 125 ms; inversion time: 2,800 ms; slice thickness: 5 mm; matrix: 512 x 512; flip angle 90 degree |
| **DWI** | repetition time: 3,000 ms; echo time: 56 ms; diffusion b-value: 1,000; slice thickness: 5 mm; matrix: 256 x 256; flip angle 90 degree |

**Supplementary figure 1.**
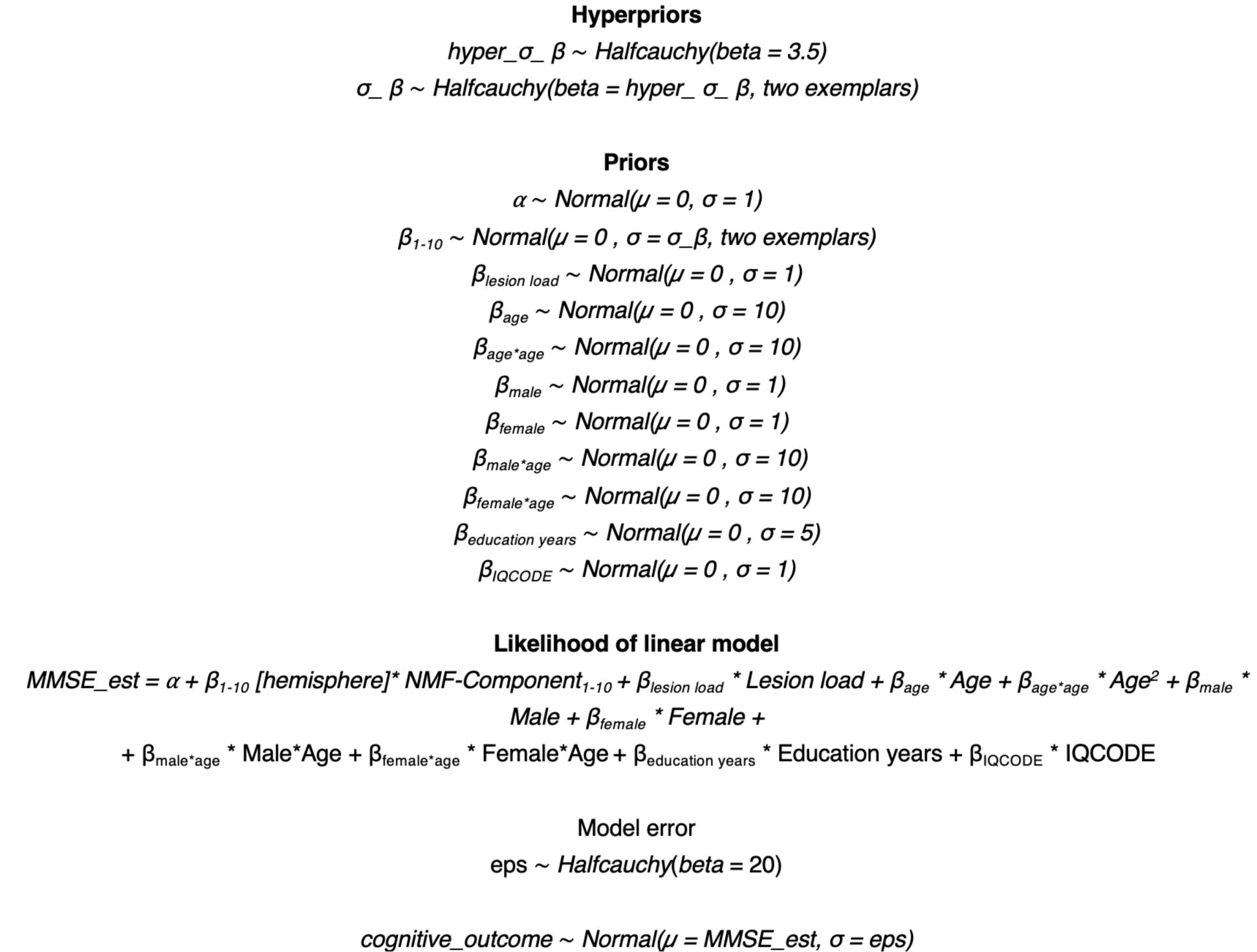
Full Bayesian model specification, exemplarily for MMSE.

